# SLIPMAT: a pipeline for extracting tissue-specific spectral profiles from ^1^H MR spectroscopic imaging data

**DOI:** 10.1101/2022.11.15.516599

**Authors:** Olivia Vella, Andrew P. Bagshaw, Martin Wilson

## Abstract

^1^H Magnetic Resonance Spectroscopy (MRS) is an important non-invasive tool for measuring brain metabolism, with numerous applications in the neuroscientific and clinical domains. In this work we present a new analysis pipeline (SLIPMAT), designed to extract high-quality, tissue-specific, spectral profiles from MR spectroscopic imaging data (MRSI). Spectral decomposition is combined with spatially dependant frequency and phase correction to yield high SNR white and grey matter spectra without partial-volume contamination. A subsequent series of spectral processing steps are applied to reduce unwanted spectral variation, such as baseline correction and linewidth matching, before direct spectral analysis with machine learning and traditional statistical methods. The method is validated using a 2D semi-LASER MRSI sequence, with a 5-minute duration, from data acquired in triplicate across 8 healthy participants. Reliable spectral profiles are confirmed with principal component analysis, revealing the importance of total-choline and scyllo-inositol levels in distinguishing between individuals – in good agreement with our previous work. Furthermore, since the method allows the simultaneous measurement of metabolites in grey and white matter, we show the strong discriminative value of these metabolites in both tissue types for the first time. In conclusion, we present a novel and time efficient MRSI acquisition and processing pipeline, capable of detecting reliable neuro-metabolic differences between healthy individuals, and suitable for the sensitive neurometabolic profiling of in-vivo brain tissue.

## Introduction

Proton (^1^H) Magnetic Resonance Spectroscopy (MRS) is an established technique for the non-invasive measurement of metabolism, offering unique insight into brain function and health. While widely used clinically for characterisation of healthy and pathological brain tissue (Oz et al., 2014), MRS can also be used to investigate functional changes in brain metabolites. Using MRS, the levels of primary excitatory and inhibitory neurotransmitters: glutamate and GABA, have been shown to modulate in response to motor learning tasks (Kolasinski et al., 2019), visual stimulation (Bednařík et al., 2015) and other experimental paradigms (Mullins, 2018) – demonstrating direct links between neurotransmitter levels; brain activation and plasticity. Glutamate responses have also been recently shown to distinguish between perceived and unperceived visual stimuli in primary visual cortex (DiNuzzo et al., 2022), crucially in the absence of neurovascular changes measured with fMRI. The specific role and action of other MRS-detectable molecules are less well understood but equally implicated in cognitive processes. For example, N-Acetylaspartic acid (NAA), one of the most abundant metabolites in the human brain, has been shown to relate to fluid intelligence and creativity (Jung et al., 2009; Nikolaidis et al., 2017).

Whilst short term alterations in metabolites such as glutamate, GABA and lactate are reasonably well characterised in response to basic visual stimuli and motor tasks over periods of minutes, longer term changes are only beginning to be established. Recent work suggests that glutamate in the lateral prefrontal cortex accumulates over period of hours in response to hard cognitive work (Wiehler et al., 2022), highlighting an important relationship between cognitive fatigue; sleep and neuro-metabolism. Alternatively, we may query which aspects of neuro-metabolism are reliably different between individuals over a longer time-period of months – with recent work demonstrating the discriminative value of choline-containing metabolites and scyllo-inositol (Wu et al., 2022).

Despite the historical and recent scientific insights derived from MRS measures of neuro-metabolism, the technique remains relatively underused in the field of cognitive neuroscience – particularly when compared to other MR-based methods such as functional-MRI and diffusion-MRI. Difficulties in obtaining high data quality; accurate signal localisation and robust analysis methods likely explain this discrepancy. However the wider availability of modern methods established by the MRS research community is expected to address these technical challenges (Bogner et al., 2021; Maudsley et al., 2021; Near et al., 2021; Öz et al., 2021; Vidya Shankar et al., 2019; Wilson et al., 2019).

In this work, we present a novel analysis pipeline designed to ameliorate several technical challenges associated with obtaining high quality neuro-metabolic spectral profiles from MRS. The most common MRS acquisition technique (single voxel MRS) is known to suffer from partial volume effects, making it difficult to avoid spectral contamination from unwanted tissue types. For instance, a 2 cm sided voxel placed in cortical grey matter will always contain a degree of white matter tissue, and variations in cortical thickness will introduce unwanted variance in the spectral data due to the metabolic differences between the two tissue types. To address this limitation, we employ a 2D semi-LASER magnetic resonance spectroscopic imaging (MRSI) sequence to acquire spectral data from an axial slice just above the corpus collosum. These spectra, containing a mixture of white and grey matter contributions, are first aligned in terms of phase and frequency (Wilson, 2019), before applying spectral decomposition (Goryawala et al., 2018) to obtain “pure” white and grey matter spectra with high SNR and spectral resolution. A series of spectral processing steps are then applied to minimise the expected spectral variations related to experimental, rather than metabolic, variability. Finally, the processed white and grey matter spectra may be investigated separately, or in combination, to form a neuro-metabolic spectral profile – amenable for further analysis with machine-learning or traditional statistical approaches. We refer to this new analysis pipeline as SLIPMAT: SpectraL Image Processing for Metabolic Analysis of Tissue-types, since metabolically distinct tissue types are extracted from a single MRSI dataset. The method is applied to MRSI data and shown to extract high quality neuro-metabolic spectral profiles, capable of distinguishing between individuals when combined with machine learning – in good agreement with our previous work (Wu et al., 2022).

## Methods

### SLIPMAT algorithm

The objective of the SLIPMAT method is to extract high quality white and grey matter spectra for analysis with machine-learning or conventional techniques. In practice, this requires MRSI and volumetric MRI datasets to be acquired from the same participant, ensuring the two datasets are spatially consistent. The simplest approach, as used here, is to ensure both scans are acquired in the same session and the participant’s head remains static throughout the session.

#### Step 1: intra-scan spectral correction and tissue decomposition

The first analysis step involves extracting the useful region of interest (ROI) in the MRSI dataset, for instance excluding voxels outside the pre-localised volume of interest (VOI) region or brain. A reference spectrum is then defined for frequency and phase alignment. Here, we used a single spectrum central to the acquisition region to minimise the potential for scalp lipid contamination. However, in more challenging brain regions, an “ideal” reference spectra could be simulated to match the acquisition protocol to ensure perfect baseline, phasing, lineshape and SNR. The frequency and phase of all other spectra within the ROI are adjusted to match the reference using the RATS method (Wilson, 2019). Following alignment, spectra are intensity normalised based on the sum of the metabolite-rich spectral data points between 4 and 0.2 ppm to reduce the influence of B1 inhomogeneity. Since broad signal components, such as residual water tails, can heavily influence normalisation, the Asymmetric Least Squares (ALS) baseline correction method (Eilers and Boelens, 2005) is applied prior to summation. Note, the baseline corrected spectra are only used to derive the normalisation scaling values, which are subsequently applied to the uncorrected spectra.

In parallel, the volumetric MRI is segmented into tissue types using standard tools (Zhang et al., 2001), before being spatially co-registered to the MRSI data – assuming no movement between the MRI and MRSI scans. Knowledge of the proportion of white and grey matter contributing to each voxel allows the spectra of the two tissue types to be determined with simple linear algebra (Goryawala et al., 2018) – eliminating partial volume effects and significantly boosting the spectral SNR relative to individual MRSI voxels. These processing steps are summarised graphically in Fig. 1.

**Fig. 1.**
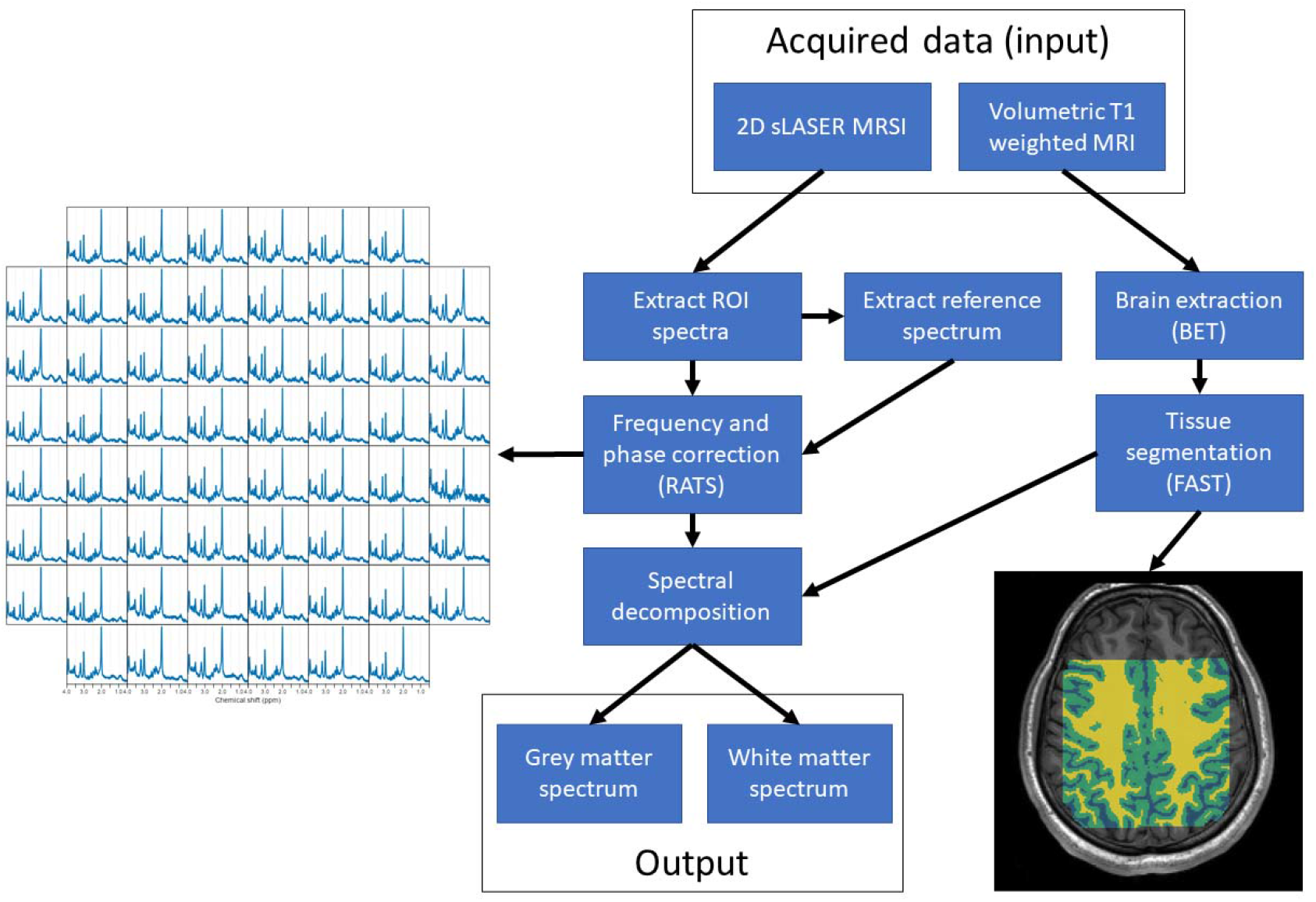
The initial phase of the SLIPMAT method, resulting in a grey and white matter spectral pair for further processing.

At this stage of the algorithm, two high quality tissue-specific spectra have been derived from the MRSI data and could be analysed with conventional fitting approaches (Near et al., 2021).

#### Step 2: inter-scan consistency correction and scaling

Once the tissue-specific spectra have been determined, a further stage of frequency and phase correction is applied (Wilson, 2019) to ensure consistency between all MRS scans involved in subsequent analysis. In our data the mean spectrum was calculated across all decomposed white and grey matter spectra and used as a reference. However, a good quality spectrum taken from a single scan, or a simulated spectrum, could also be used. A simple zero-order baseline correction step is applied, subtracting any intensity offset from zero as measured from a flat spectral region (0 to -1 ppm). A variable degree of gaussian line broadening is also applied to each spectrum to reduce the influence of inconsistent shimming between spectra. For our data, the largest FWHM of the tNAA resonance was 0.06004 ppm, therefore all spectra were broadened to 0.061 ppm.

Zero-filling to twice the original length is performed to improve spectral SNR (Bartholdi and Ernst, 1973), before discarding the imaginary component and removing spectral points outside the metabolite-rich spectral region (4 to 0.2 ppm). ALS baseline correction is applied to eliminate signals from residual water or broad out-of-volume scalp lipids. Finally, white and grey matter spectra are separately scaled by the summation of spectral data points.

These spectra, derived from the same acquisition, may be used individually or concatenated to form a single feature vector for further analysis (Fig. 2).

**Fig. 2.**
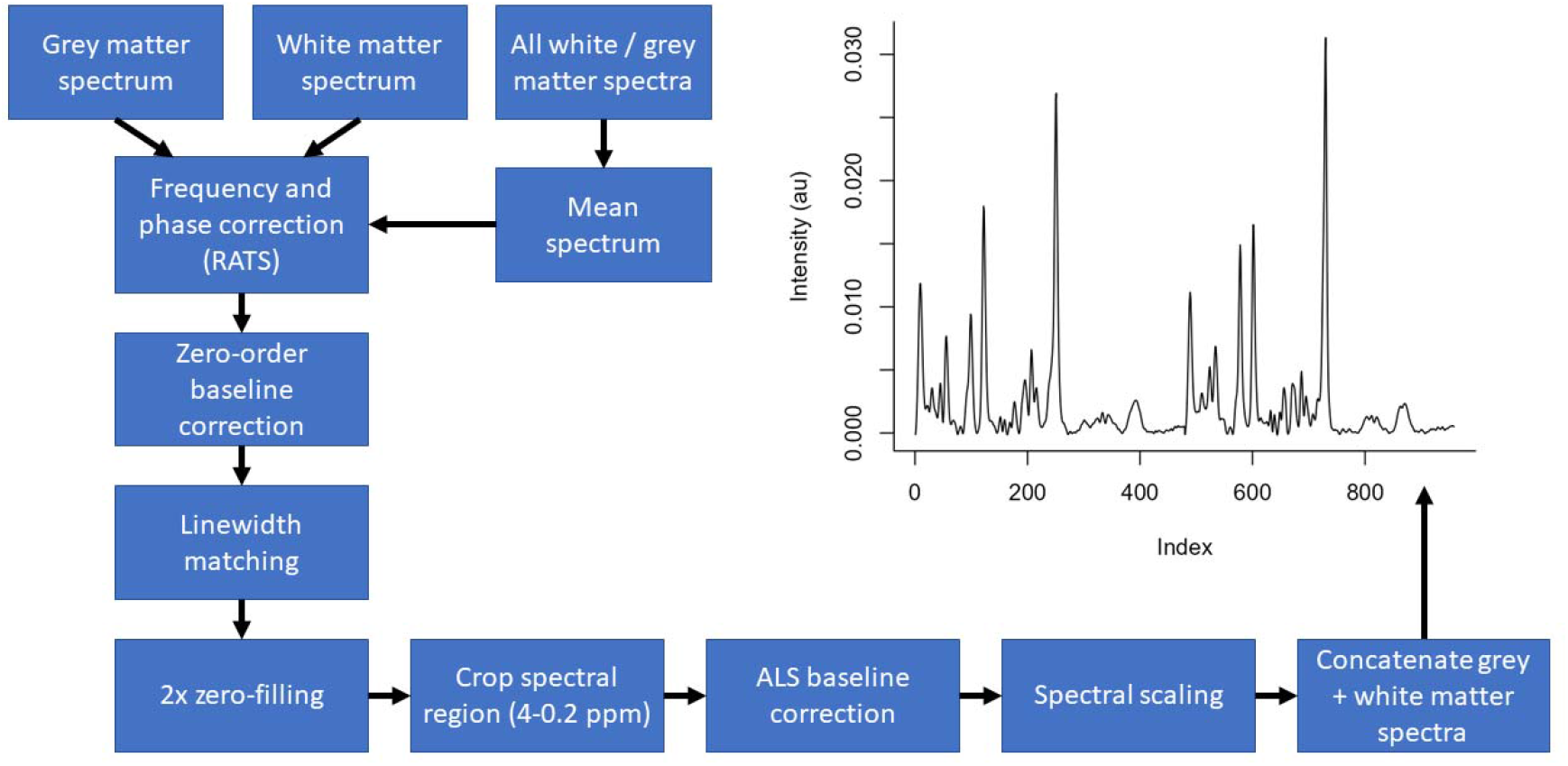
The final phase of the SLIPMAT method, designed to produce high quality spectra with consistent linewidth, baseline, phase and frequency offsets between scans.

All spectral processing and co-registration steps are implemented in the open-source spectral analysis package spant (Wilson, 2021a) and image segmentation is performed with FSL (Woolrich et al., 2009).

### MR acquisition

A validation MR dataset was acquired at the Centre for Human Brain Health from 8 healthy adults (2 female, mean age 21 years) with a 3 T Siemens Magnetom Prisma (Siemens Healthcare, Erlangen, Germany) system using a 32-channel receiver head coil-array. A T1-weighted MRI scan was acquired with a 3D-MPRAGE sequence: FOV = 208 × 256 × 256 mm, resolution = 1 × 1 × 1 mm, TE / TR = 2 ms / 2000 ms, inversion time = 880 ms, flip angle = 8 degrees, GRAPPA acceleration factor = 2, 4 minute 54 second scan duration. Water supressed MRSI data were acquired with 2D phase-encoding: FOV = 160 × 160 × 15 mm, nominal voxel resolution 10 × 10 × 15 mm, TE / TR = 40 ms / 2000 ms, complex data points = 1024, sampling frequency = 2000 Hz. The MRSI slice was aligned axially in the subcallosal plane with an approximately 1 mm gap from the upper surface of the corpus callosum. The semi-LASER method (Scheenen et al., 2008) was used localize a 100 × 100 × 15 mm VOI, central to the FOV, four saturation regions were placed around the VOI prescribing a 100 × 100 mm interior, and an additional four saturation regions were positioned to intersect the four corners of the semi-LASER VOI to provide additional scalp lipid suppression. The total acquisition time for a single MRSI scan was 5 minutes and 6 seconds, and three MRSI datasets were acquired sequentially during the same session to assess technical repeatability.

### Algorithm validation

The SLIPMAT method was applied to 60 voxels taken from the central 8 × 8 grid of each MRSI dataset, with the 4 corner voxels being excluded due to their proximity to the diagonal saturation regions. Eight participants were scanned, and three MRSI scans were acquired in each of the eight sessions, resulting in 24 pairs of processed white and grey matter spectra for further analysis. The study was conducted according to the principles expressed in the Declaration of Helsinki and participants gave written informed consent before data collection. All T1 scans were defaced (fsl_deface tool) as a first step for data sharing purposes.

To validate the approach the dataset was interrogated in three ways. Firstly, principal component analysis (PCA) was applied to the processed 48 white and grey matter spectra to explore the greatest sources of spectral variance. Secondly, the white and grey matter spectral pairs were concatenated and PCA was applied to establish the reliability and uniqueness of an individual’s neuro-metabolic profile. Finally, one-way ANOVA was applied to each spectral data point, with the participant ID used as a factor variable, to further examine which spectral regions were most unique to an individual. Results were compared with known spectral differences between white and grey matter, and a previous study exploring individual differences in neurometabolism to assess consistency.

The segmented MRI volume and MRSI voxels are spatially co-registered, based on the geometry information embedded in the exported data files (affine matrix), and therefore our method assumes that the subjects head remains static during the session. To assess the sensitivity of our method to movement, the MRSI affine matrix was manipulated to simulate shifts in the x-y plane of the scanner coordinate system. 0, 2, 5, 8, 10 and 15mm displacements were applied to each session, with the first scan remaining unaffected and the second and third shifted along the x = y line. For example, with the 2mm shift, the first scan was not modified, the second scan was shifted 1.414mm along the x and y directions and the third scan was shifted 1.414mm along the -x and -y directions. The SLIPMAT method was applied to each displacement level and PCA was performed on the concatenated white and grey matter spectral pairs (as above) to assess how spatial shifts influence the uniqueness of an individual’s neuro-metabolic profile. PCA loadings calculated from the 0mm shift were used for all displacements to ensure consistency between analyses and aiding visual comparison.

The SLIPMAT method exacts spectra corresponding to the two tissue components, grey and white matter, which are known to have distinct spectral patterns. To explore the possibility of spectral variability within these tissues types we applied spectral decomposition to three tissue classes: 1) grey matter, 2) central white matter and 3) peripheral white matter. Central and peripheral white matter regions were distinguished by calculating the median proportion of segmented white matter across the analysis region (60 voxels). Voxels with greater than 1.5 times the median white matter proportion had their white matter contribution assigned to central white matter, whereas those voxel below the threshold were assigned peripheral white matter. A cut-off of 1.5 times the median was chosen to approximately balance the SNR of the two decomposed white matter types. The SLIPMAT method was applied as previously described, however for this analysis the output was increased (from two) to three spectra per MRSI acquisition, one per tissue class. Finally, PCA was applied to the individual spectra to assess any metabolic differences between central and peripheral white matter across all the acquired MRSI data.

All MRS analysis, statistics and machine-learning was performed with the R statistical computing platform (R Core Team, 2021) and analysis scripts used to generate the figures and tables in this paper are available from https://github.com/martin3141/slipmat_paper. MRS and MRI data are available in NIfTI format (Clarke et al., 2022) from Zenodo in BIDS format: https://doi.org/10.5281/zenodo.7189139.

## Results

Compared to other MR techniques, MRS is particularly sensitive to inhomogeneities in the static field across the VOI. Spatially dependant changes in the frequency and phase of metabolite resonances results in destructive interference when combing spectra, either through simple spatial averaging or spectral decomposition. To mitigate these effects, the SLIPMAT method first corrects the spatially dependant frequency and phase inconsistencies using the RATS algorithm – designed to be robust to confounding spectral features, such as baseline instability (Wilson, 2019). To evaluate the improvement in spectral quality, the pre-processing pipeline (Fig 1) was applied with and without the frequency and phase correction (RATS) step. Fig. 3 shows how the SNR and spectral linewidth of decomposed white and grey matter spectra are significantly improved through the correction of spatial phase and frequency inconsistencies, as performed by the first phase of the pipeline (Fig 1). Average improvements in SNR across all MRSI scans were 51% (178 to 268, paired t-test p=2.2e-16) accompanied by a 20% reduction (0.056 to 0.045 ppm, paired t-test p=5.9e-5) in spectral linewidth, measured from the tNAA resonance at 2.01 ppm. These improvements are visually confirmed by comparing the grey matter lactate doublet at 1.3 ppm – which is clearly distinct from the noise in the corrected data (Fig 3C) compared to the uncorrected (Fig 3A).

**Fig. 3.**
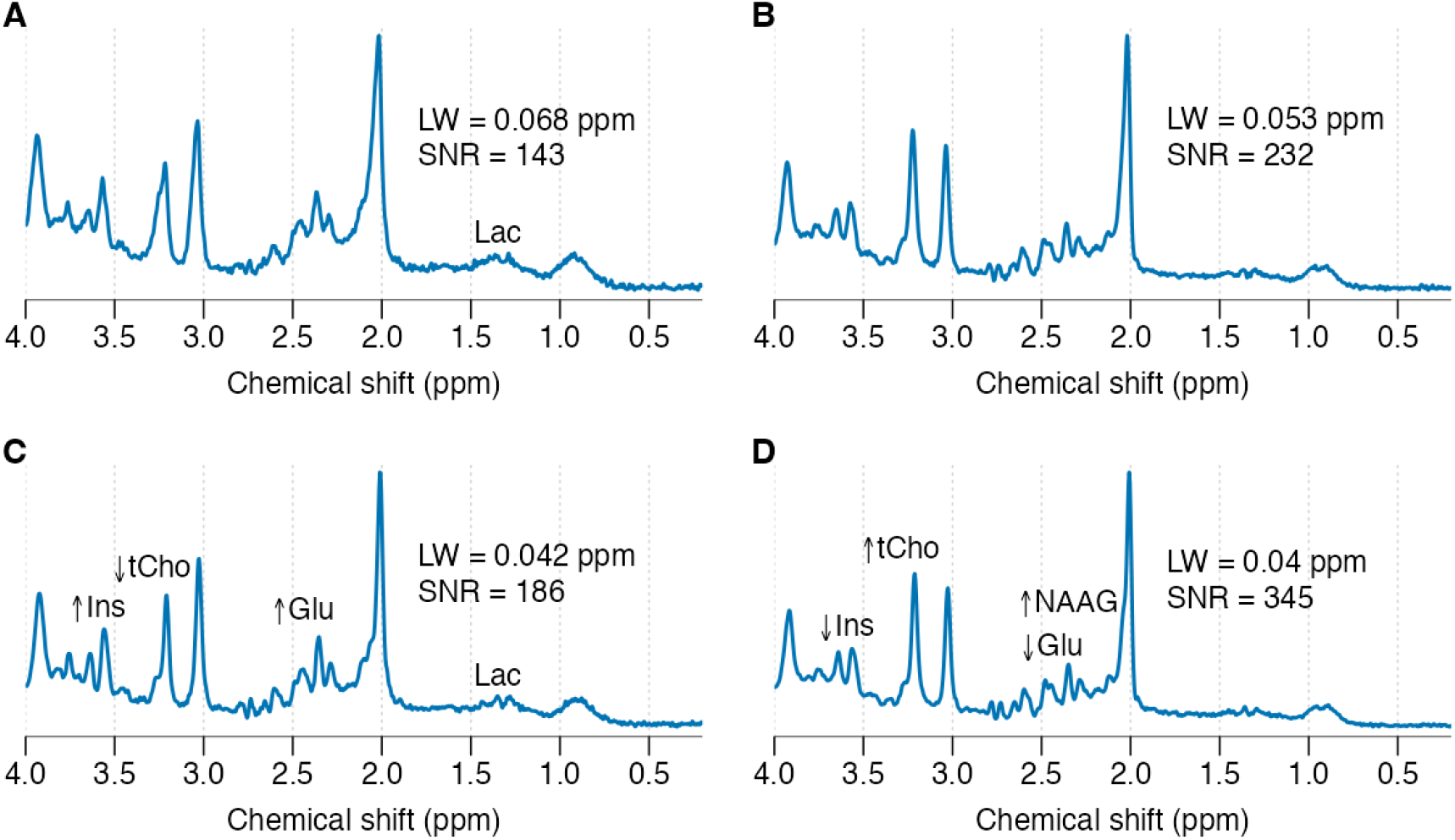
Representative comparison of spectral quality with and without the frequency and phase correction (RATS) step for decomposed grey (A, C) and white (B, D) matter spectra. Corrected spectra (C, D) show improved SNR and spectral resolution compared to uncorrected (A, B). Abbreviations: LW - linewidth, SNR - signal to noise ratio, Ins - myo-inositol, tCho – total-choline, Glu – glutamate, NAAG - N-Acetylaspartylglutamic acid, Lac – lactate.

The metabolite profiles of white and grey matter are known to be different (Pouwels and Frahm, 1998) and our results support these observations. For example, glutamate and myo-inositol are stronger in parietal grey matter, whereas NAAG and total-choline are stronger in parietal white matter (Fig. 3 C vs D).

PCA was applied to the full set of white and grey matter spectra across 8 individuals to confirm the primary sources of spectral variance arise from genuine neuro-metabolic differences, rather than spectral artifacts such as residual water signals and scalp lipid interference. Fig. 4 shows 80% of the variance (PC1) can be explained by differences between the two tissue types. PCA loading plots show which spectral regions contribute most strongly to the scores, with the loadings plot for PC1 (Fig 4. part B) highlighting total-choline (3.2 ppm), NAAG (2.0 ppm) and glutamate (2.4 ppm). The intensity of these resonances agrees with expected differences between white and grey matter, with total-choline and NAAG being elevated in white matter (positive scores in PC1) and glutatmate being elevated in grey matter (negative scores in PC1).

**Fig. 4.**
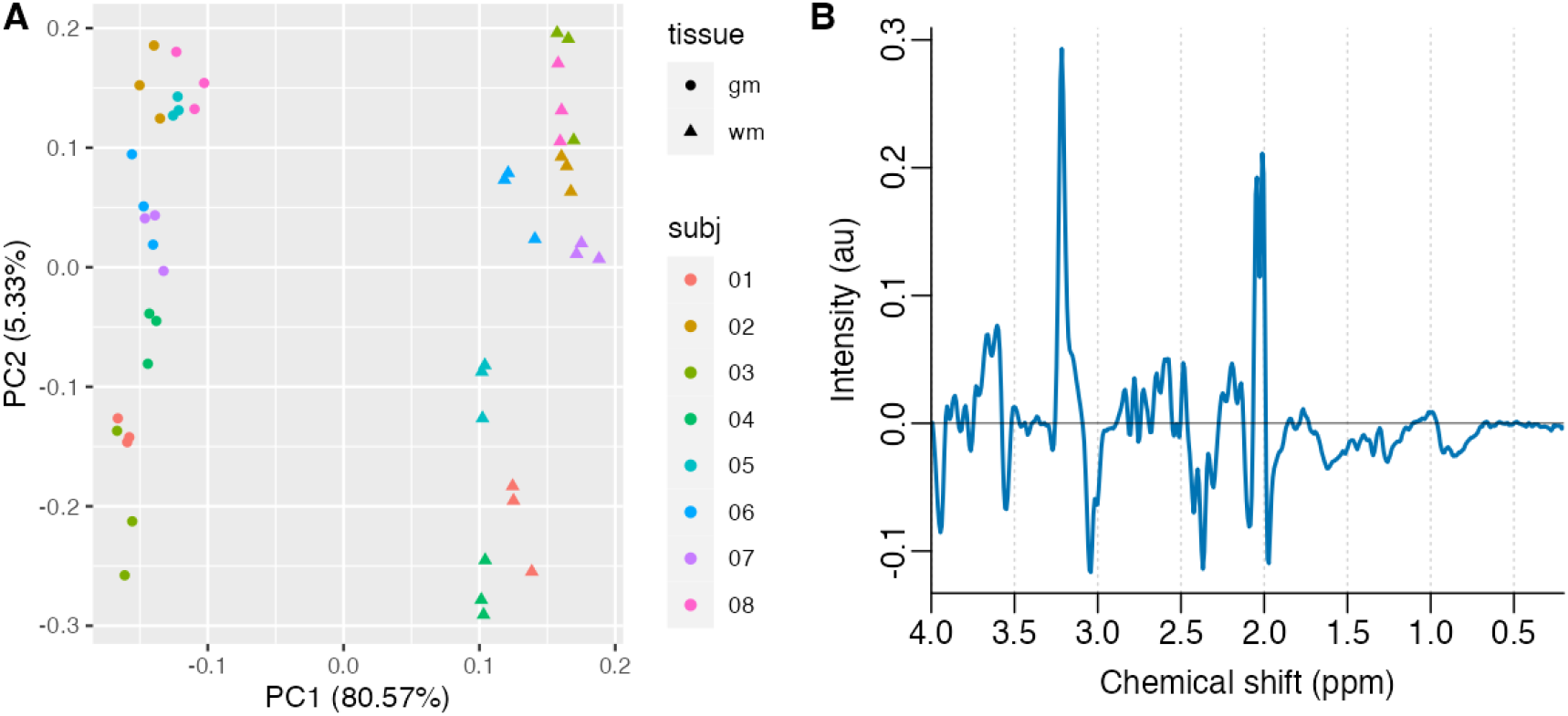
PCA scores (A) and loadings for PC1 (B) of all white and grey matter spectra following processing with SLIPMAT. Clear separation is seen between the two tissue types in agreement with known metabolic characteristics.

In addition to tissue-types, Fig. 4 also demonstrates consistent spectral differences between subjects, with some clustering of repeat measures observed in PC2. To explore these differences further, the grey and white matter spectra for each MRSI dataset were combined into a single feature vector – allowing both tissue types to contribute to an individual’s neuro-metabolic spectral profile. PCA scores of these combined profiles (Fig. 5 A) shows a clear separation between subjects – with the variance between repeats (technical variance) being smaller than the variance between individuals (biological variance). Inspection of the PCA loadings (Fig. 5 B-E) highlights the importance of total-choline (3.21 ppm) and scyllo-inositol (3.34 ppm) in discriminating between individuals, in strong agreement with our previous work (Wu et al., 2022).

**Fig. 5.**
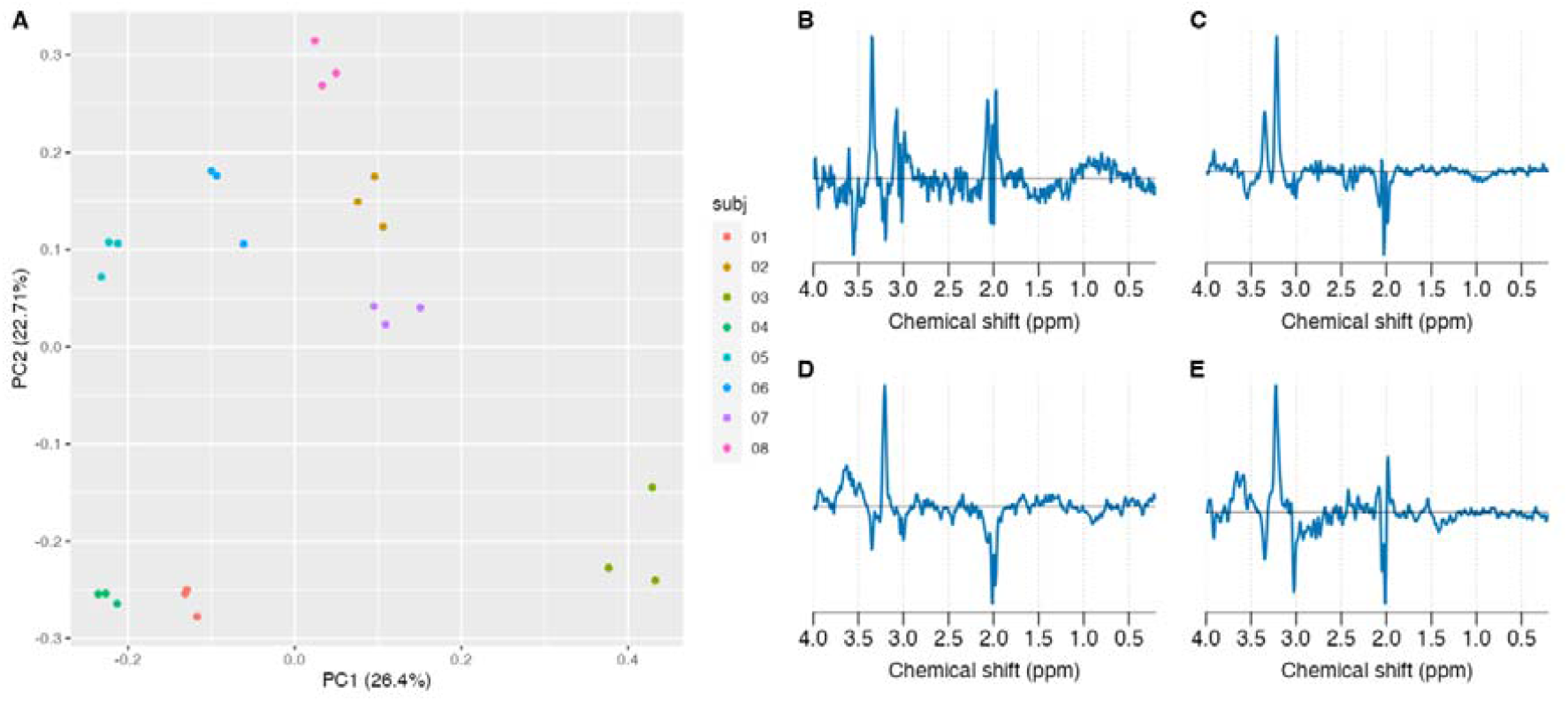
PCA scores (A) and loadings (B-E) of concatenated white and grey matter spectral pairs following processing with SLIPMAT. Loadings for the PCs have been split into white and grey matter and displayed separately to aid spectral interpretation. Grey matter spectral scores are shown in parts B) and D) corresponding to PC 1 and PC 2 respectively. White matter spectral scores are shown in parts C) and E) corresponding to PC 1 and PC 2 respectively.

Unsupervised methods such as PCA are informative for validation purposes, but in practice the neuro-metabolic influence of a known intervention or disease state is more commonly sought. One simple approach is to apply a conventional statistical technique, such as a t-test in the case of two groups, to each spectral data-point to identify key frequencies and metabolites related to the known grouping. Here we use a one-way analysis of variance (ANOVA) to assess which spectral features are the most discriminatory between individuals for both white and grey matter.

Strong individual differences in total-choline and scyllo-inositol are apparent from the high ANOVA F-statistics in those spectral regions (Fig. 6), with larger differences observed in white matter compared to grey matter. We note the strong similarities between Fig. 6 B) in this report and Fig. 3 from (Wu et al., 2022), with both plots highlighting similar spectral regions, despite differences in: scanner hardware and location; participant cohort; acquisition protocol (single-voxel PRESS vs semi-LASER MRSI) and analysis pipeline (Wilson, 2021b). We also note, for the first time, that total-choline and scyllo-inositol are strong discriminatory features in white matter, whereas this was not possible to determine from our previous work, where a single-voxel was placed in the ACC region containing an average tissue proportion of 75% grey matter and 25% white matter.

**Fig. 6.**
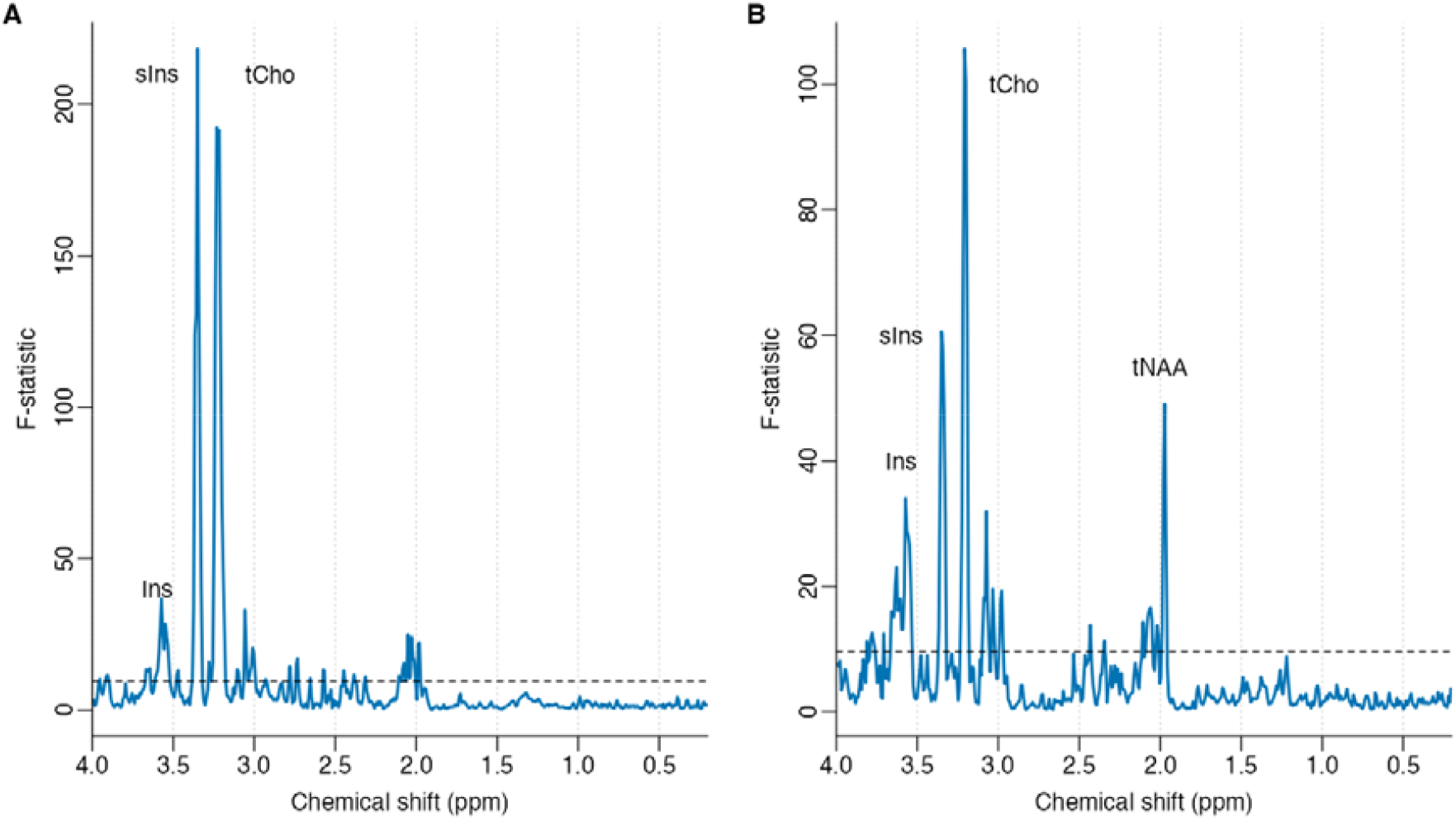
Spectral ANOVA results performed on each frequency domain point independently and grouped over participants for decomposed (A) white and (B) grey matter spectra. Large F-statistic values correspond to a strong difference in the spectral intensity between participants. Dashed horizonal lines represent the Bonferroni corrected significance threshold (p < 1e-4). Predominant spectral differences between participants are clear in both tissue types at 3.35 and 3.2 ppm assigned as sIns and tCho respectively.

The results from simulated spatial displacements between the MRSI and anatomical scan are presented in Fig. 7. Only minor changes in PCA scores are observed for displacements up to 10mm, demonstrating that SLIPMAT is tolerant to displacements below this level.

**Fig. 7.**
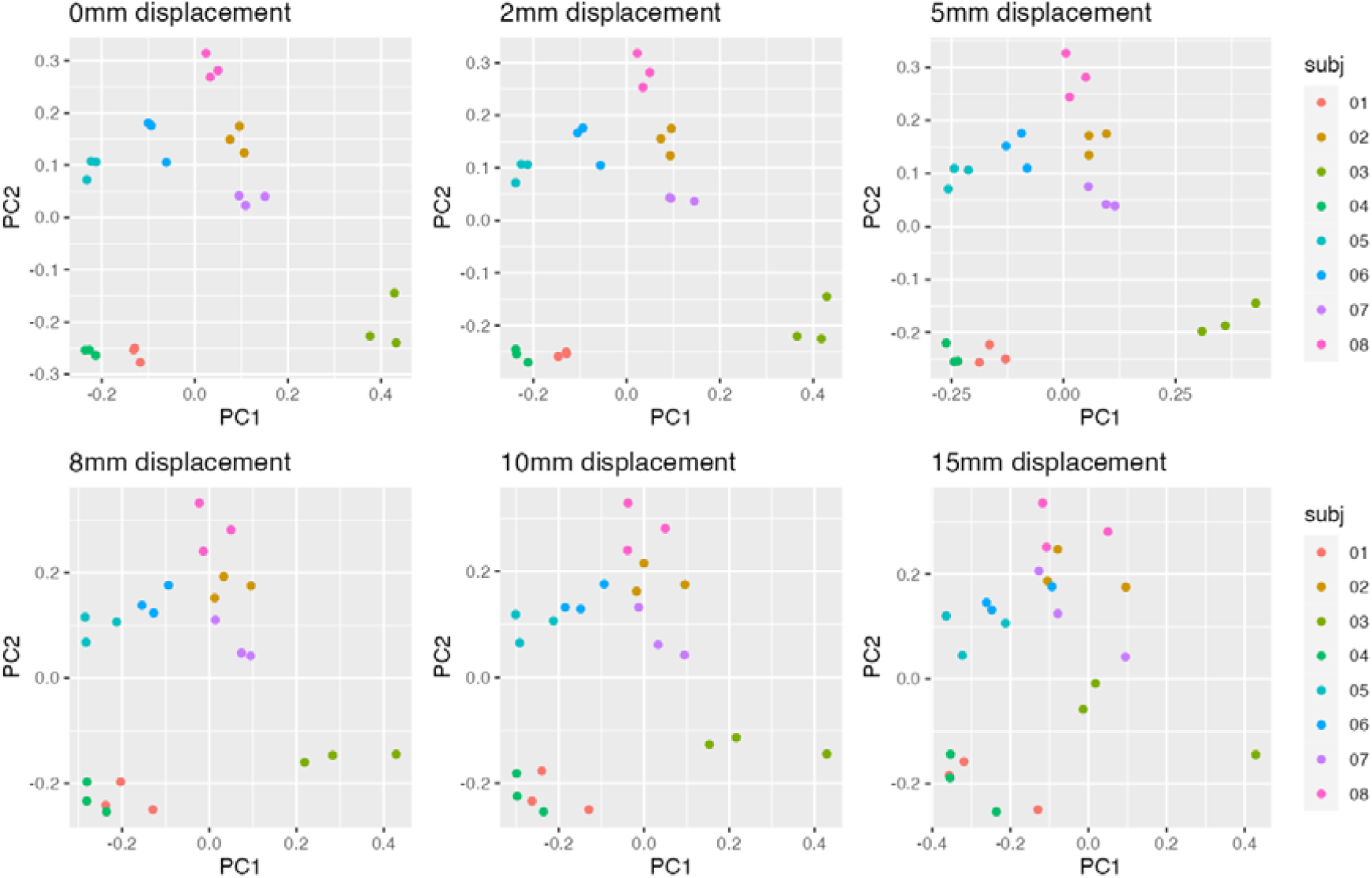
PCA scores of concatenated white and grey matter spectral pairs following processing with SLIPMAT. Increasing levels of spatial displacement have been artificially applied in each panel to explore how movement between anatomical and MRSI acquisitions is likely to degrade the uniqueness of an individual’s neurometabolic spectral profile.

Larger displacements cause the triplicate scans, belonging to each subject, to become less tightly clustered and increase their overlap with other subjects – indicating the spectral variability from spatial displacement is becoming greater than the spectral variability between participants.

The SLIPMAT method assumes MRSI data may be split into two primary tissue types, white and grey matter, and these are chosen due to their known distinct spectral features (Fig. 4). However, less is known about how metabolite levels vary spatially within these tissue types, therefore we extended the SLIPMAT method to examine differences in central and peripheral white matter regions. Fig. 8 shows how the largest source of variance (PC1, 73%) remains the difference between white and grey matter, however splitting central and peripheral white matter shows smaller (PC2, 9.7%), but consistent, differences between these tissue types. For each subject, central white matter regions show higher scores in PC2 relative to peripheral white matter, corresponding to greater levels of tCho and lower levels of tNAA (Fig. 8 part B). Notably, the variance between central and peripheral white matter (PC 2) is orthogonal to the variance between white and grey matter (PC 1), suggesting the differences between central and peripheral white matter cannot be explained by differing levels of erroneous spectral contamination between white and grey matter. If spectral contamination between white and grey matter was a primary factor, scores for peripheral white matter would be expected to move closer to the grey matter cluster, e.g. negative scores for PC1.

**Fig. 8.**
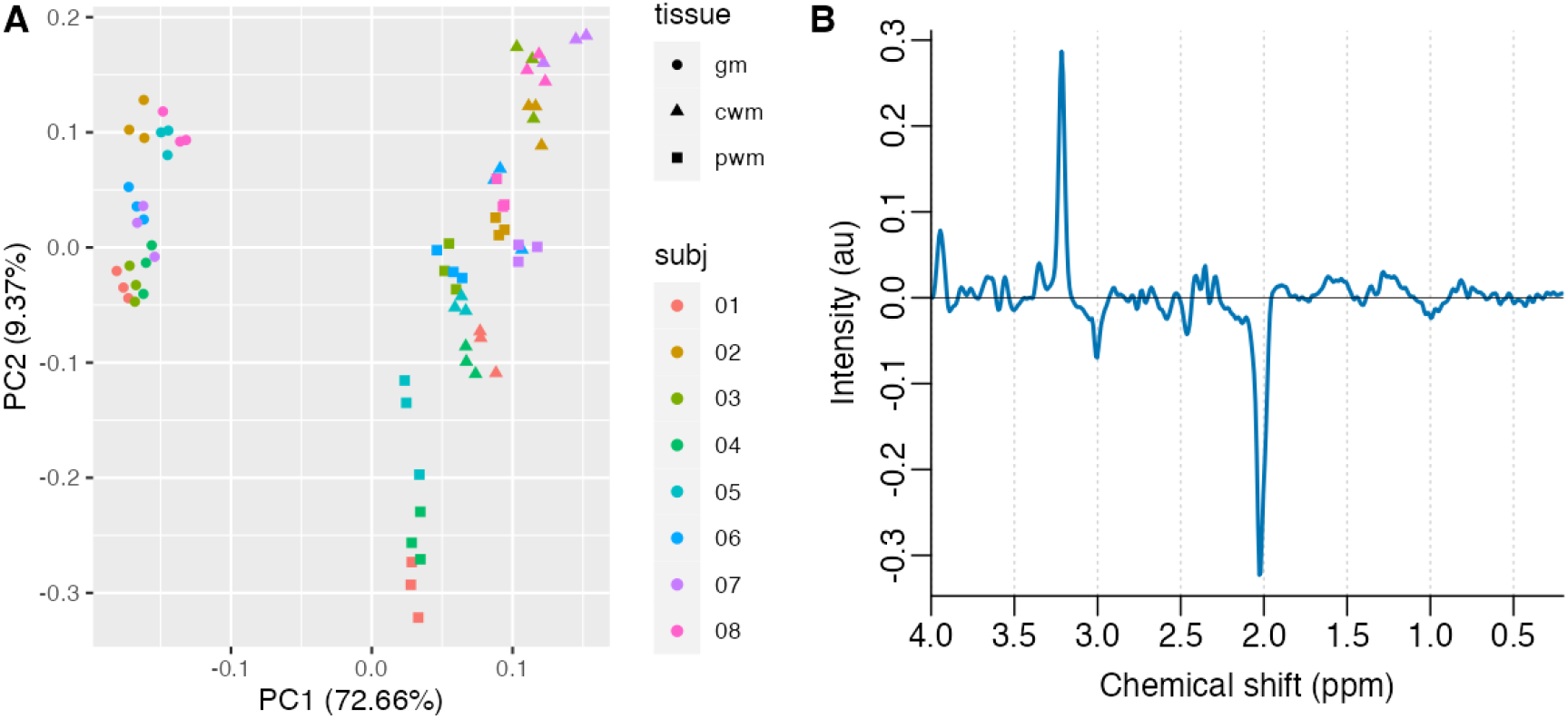
PCA scores (A) and loadings for PC2 (B) of a three-way decomposition of all grey matter (gm), central white matter (cwm) and peripheral white matter (pwm) spectra following processing with SLIPMAT. Clear separation is seen between grey and white matter (PC1), with additional and orthogonal variability apparent between central and peripheral white matter (PC2).

## Discussion

The primary goal of MRS analysis is to extract biologically meaningful measures, whilst reducing the influence of variability from technical / experimental factors. In this work, we demonstrate how the sequential application of discrete spectral processing steps can achieve this aim. Partial volume effects; frequency and phase inconsistencies; baseline; and linewidth variability are reduced using a variety of techniques (Eilers and Boelens, 2005; Goryawala et al., 2018; Wilson, 2019), and combined into a novel and fully automated analysis pipeline: SLIPMAT. The method is validated using MRSI data from 8 healthy participants, highlighting subtle spectral features associated with individual differences in neuro-metabolism in strong agreement with previous work (Wu et al., 2022).

Conventional MRS analysis relies heavily on parametric least-squares fitting of a known basis-set of metabolite, macromolecule and lipid signals (Near et al., 2021). However, in this work, we take the approach of extracting spectral “fingerprints” for further analysis with multivariate and machine learning techniques, a strategy more commonly associated with analytical chemistry – known as chemometrics (Wehrens, 2020). One significant advantage of this approach includes the potential to discover spectral features that are not typically incorporated into a fitting basis-set, e.g., novel metabolites. In addition, this methodology is relatively simple in design compared to modern MRS fitting algorithms, where differing algorithmic approaches result in weak agreement between software packages (Craven et al., 2022). A modular, pipeline-based, design simplifies the evaluation of each individual processing step, allowing improvements to be made in a more systematic and transparent fashion. For instance, although not required for our data, an additional preprocessing step to incorporate scalp-lipid suppression could be incorporated for more technically challenging acquisition protocols such as 3D-FID-MRSI (Bilgic et al., 2014; Moser et al., 2020).

To the best of the author’s knowledge, this study is the first to investigate the efficacy of the spectral decomposition method for conventional 2D semi-LASER MRSI, demonstrating excellent spectral resolution and SNR (Fig. 3) when combined with spatial frequency and phase correction (Wilson, 2019). Whilst 2D semi-LASER is limited to a single slice, the 5-minute acquisition protocol is time efficient and comparable to a clinical single voxel duration.

In this study, we incorporate the spectral decomposition method into our processing pipeline (Goryawala et al., 2018), however we note that the SLIM family of localisation methods could also be used in its place (Hu et al., 1988; Lee et al., 2017). Both approaches make use of high-resolution segmented imaging data to extract spectra associated with each segmented tissue type from an MRSI acquisition. Example white and grey matter spectral output from the first phase of the SLIPMAT method (Fig. 3C, D) show good visual agreement with spectra obtained with BASE-SLIM (Adany et al., 2016) suggesting that both methods are likely to yield similar results. However, BASE-SLIM requires the additional acquisition of B0 and B1 maps, which are subsequently incorporated into the reconstruction, whereas in this work we combine frequency and phase correction (Wilson, 2019) with spectral decomposition as separate processing steps, without requiring an additional B0 acquisition. To the best of the author’s knowledge, a direct comparison between SLIM based approaches and spectral decomposition has not been reported and would therefore make an interesting future study.

We tested and validated our proposed method using 2D MRSI acquired from the centrum semiovale in healthy participants and assumed that tissue metabolism in this region could be well characterized into two types: white and grey matter. However, additional variability in metabolite levels is likely to exist within these tissue types, and we used an exploratory three compartment model to show that peripheral and central white matter have small, but consistent, differences in tNAA and tCho (Fig. 8). These findings are not surprising, as low level spatial heterogeneity in metabolite levels have also be observed with high resolution MRSI at 7T (Hangel et al., 2018). Additional spatial variability is therefore expected for the analysis of greater brain volumes, for example whole-brain MRSI, and a more detailed classification of tissue types may be required. However, this represents a trade-off, where the addition of more tissue types and regions of interest will reduce spectral SNR, potentially obscuring low level metabolites that could act as novel diagnostic markers. Furthermore, the use of SLIPMAT for focal disease, such as brain tumours or stroke, may not be optimal due to the loss of spatial information. For these cases, one potential extension to the technique could involve modelling the individual (spatially resolved) MRSI spectra as a linear combination of the “pure” white and grey matter spectra derived from the SLIPMAT method. Local discrepancies from the two-compartment model could be identified by evaluating the fitting residuals from this process, potentially defining additional tissue classes that could be automatically incorporated into a subsequent SLIPMAT analysis with a higher number of tissue components. We also note that low-rank decomposition methods, such as SPICE (Lam and Liang, 2014; Ma et al., 2015), may be more appropriate for the analysis of MRSI with multiple unknown tissue compartments.

The importance of correcting spatially dependant frequency shifts in MRSI data processing have been shown previously, with alignment methods including cross-correlation (Ebel and Maudsley, 2003), water reference data (Maudsley et al., 2006) and magnitude-based spectral correction (Le Fur and Cozzone, 2014). Here we use the RATS method (Wilson, 2019) – originally developed for correcting frequency and phase inconsistencies for single-voxel MRS. The method is shown to perform well for MRSI data (Fig. 3) and may be especially suited to this application due to its robustness to baseline instability – a common artefact for MRSI datasets. More generally, this work supports the use of correcting spatially dependant frequency and phase differences in MRSI to improve data quality, in agreement with recent recommendations (Maudsley et al., 2021).

Single-voxel MRS remains more popular than MRSI due to reduced technical demands (Wilson et al., 2019), however we envisage our proposed method may be an attractive alternative for certain applications. In clinical MRS, reduced scan time is particularly crucial, and the opportunity to acquire high quality white and grey matter spectra, free from partial volume effects, within the same 5-minute scan represents a compelling advance. Potential clinical applications include biomarker discovery for early-stage neurodegeneration or mental illness – where changes in conventional MRI are either absent or occur too late in the disease process for effective treatment. In cognitive neuroscience, where relevant changes in neurometabolism are likely to be small, partial volume effects introduce unwanted variability and have the potential to dominate over genuine metabolic modulations. We anticipate the proposed method will be a more sensitive tool for detecting small changes in metabolite levels due to the elimination of these partial volume effects with spectral decomposition.

Whilst we chose a chemometric approach for spectral processing, we note that conventional fitting methods could also be applied directly to the decomposed white and grey-matter spectra (Fig. 1). This has been demonstrated previously with a high-resolution whole-brain MRSI protocol (Goryawala et al., 2018). A dedicated comparison between chemometric and conventional fitting approaches is warranted for larger MRS datasets, and we anticipate this will be aided by recent effects to lower barriers associated with MRS data sharing (Clarke et al., 2022; Soher et al., 2022).

One limitation of the study is that subjects were scanned in triplicate in the same session, and therefore the observed differences could be related to session, rather than biological, variability. Comparing results from this study with our previous work, where subjects were scanned using SVS across multiple sessions, we note the strong similarities between Fig. 6B) from this report and Fig. 3 from (Wu et al., 2022) which both show total-choline and scyllo-inositol as being strong discriminatory features between healthy individuals. Considering the large effect sizes detected in both studies, we believe it is more likely that biological, rather than session dependant, factors underly the observed discriminatory spectral features.

Movement between, and during, the anatomical and MRSI scans has the potential to bias metabolite profiles. Spatial shifts were artificially applied to the MRSI data to estimate the sensitivity of SLIPMAT to the influence of typical head movements, and the method was shown to be robust to displacements up to approximately 10mm (Fig. 7). An assessment of typical head movements in MRI was conducted using 42,874 scans from the UK Biobank dataset (Alfaro-Almagro et al., 2018), showing that 99% of participants moved less than 5mm over a 10 minute window (Hess et al., 2022). In this study, the anatomical and MRSI scan were both acquired within 10 minutes, suggesting that head movement is unlikely to be significant problem for our proposed protocol and analysis pipeline. For cases where higher levels of movement are anticipated, for example paediatric studies, the post-acquisition alignment of MRSI and anatomical imaging could be performed using a rigid registration method (Maudsley et al., 2006) to maintain spatial consistency between the scans.

Potential disadvantages of the proposed method include difficulties in combining datasets acquired under different experimental conditions. For example, a dataset acquired at 1.5 and 3 T magnetic field strengths would have differing spectral features for any metabolites containing J-coupled resonances, and these features may dominate over biological variability. Fitting based methods are expected to be less susceptible to these differences, since the basis set may be adapted to match the magnetic field strength and specific acquisition protocol (Near et al., 2013). Future work could include the evaluation of more modern machine learning approaches, such as deep learning, to reduce the impact of combining spectral profiles acquired with differing acquisition protocols.

In conclusion, we have developed a novel and time efficient MRSI acquisition and processing pipeline, capable of detecting reliable neuro-metabolic differences between healthy individuals. Future work will investigate how rapid tissue-specific neuro-metabolic profiling can be applied to neuroscientific investigation and clinical studies of the brain.

## CRediT authorship contribution statement

Olivia Vella: Validation, Formal analysis, Investigation, Visualisation. Andrew P. Bagshaw: Conceptualisation, Writing – Review & Editing. Martin Wilson: Conceptualisation, Methodology, Software, Validation, Formal analysis, Investigation, Data curation, Writing – Original Draft, Visualisation, Supervision.

## Data and code availability

All MR data is available from Zenodo in BIDS format, doi: 10.5281/zenodo.7189140. Code used to generate the results is available from GitHub, https://github.com/martin3141/slipmat_paper.

